# DESeq2-MultiBatch: Batch Correction for Multi-Factorial RNA-seq Experiments

**DOI:** 10.1101/2025.04.20.649392

**Authors:** Julien Roy, Adrian S. Monthony, Davoud Torkamaneh

## Abstract

RNA sequencing (RNA-seq) experiments frequently encounter batch effects that can significantly distort biological interpretations, particularly in complex, multifactorial studies where biological variables interact with experimental batch conditions. Existing batch correction tools primarily address technical variability and often neglect these critical interaction effects, resulting in incomplete adjustments. To address this gap, we introduce DESeq2-MultiBatch, a novel, lightweight batch correction method implemented entirely within the DESeq2 analytical framework. Unlike conventional approaches, DESeq2-MultiBatch directly leverages DESeq2’s internal model estimates to correct raw gene count data, accurately adjusting for experimental batch effects, including interactions with biological variables. Here, we demonstrate that DESeq2-MultiBatch effectively mitigates batch-related variability while preserving genuine biological differences. Benchmarking against widely used methods highlights DESeq2-MultiBatch as a robust, practical solution for improving exploratory data visualization and downstream analyses in multifactorial RNA-seq studies.

**Availability and implementation:** https://github.com/julienroyulaval/DESeq2-MultiBatch

**Supplementary information:** Supplementary data are available at Bioinformatics online.

## Introduction

RNA sequencing (RNA-seq) has emerged as an indispensable tool in modern biology, enabling detailed analyses of gene regulation, cellular processes, and organismal responses under diverse biological conditions (Stark, Grzelak and Hadfield 2019). Various computational frameworks have been developed to process RNA-seq data, each employing different statistical and bioinformatics strategies (Costa-Silva et al. 2021). However, RNA-seq experiments frequently encounter batch effects, defined as sub-groups of measurements showing qualitatively different behavior across conditions due to unwanted technical variability or non-biological differences introduced during experimental preparation, sequencing, or data processing (Goh, Wang and Wong 2017, Leek et al. 2010). Although careful experimental design can mitigate these effects, practical constraints frequently necessitate multi-batch studies, making complete avoidance challenging (Van Den Berge et al., 2019; Ye et al., 2023).

To date, several computational strategies have been developed to address batch effects, primarily by integrating a batch factor into models of gene expression. Simple linear models, such as mean-scaling and zero-centering, fall under this class (Goh, Wang and Wong 2017). The empirical Bayes algorithm (e.g., ComBat) was among the first widely-adopted methods, effectively normalizing additive and multiplicative batch effects (Johnson et al. 2007). Limma, a popular package for gene expression analysis, incorporates batch-effect removal directly into its linear model framework, streamlining analyses (Ritchie et al. 2015). Recently, ComBat-seq extended the original ComBat algorithm specifically for RNA-seq data, using negative binomial regression to better handle count data typical of RNA-seq experiments (Zhang, Parmigiani and Johnson 2020). Additionally, surrogate variable analysis (SVA) offers another approach by identifying part of the data associated with batch variation, estimating the effect with singular value decomposition and removing it from data via regression (Leek 2014, Leek and Storey 2007). Most recently, machine learning-based approaches have also been integrated into batch-effect detection and correction workflows, demonstrating promising results in complex scenarios (Sprang, Andrade-Navarro and Fontaine 2022).

Performance assessments of existing batch correction methods, conducted with both real and simulated datasets, highlight their effectiveness as highly context-dependent (Chen et al. 2011, Liu and Markatou 2016). Although many algorithms are robust to moderately confounded experimental designs, their effectiveness diminishes significantly when biological factors and batches are highly confounded, potentially leading to the loss of true biological signals (Molania et al. 2023, Zhou, Chi-Hau Sue and Bin Goh 2019). Furthermore, batch-correction techniques modifying raw count data require careful application, as they can inadvertently mask genuine biological differences, particularly in unbalanced experimental designs (Nygaard, Rødland and Hovig 2016).

Most existing batch correction evaluations have utilized medical or clinical datasets, thus limiting their applicability to non-model species (i.e., plants), where broader experimental designs are commonplace. RNA-seq studies in plants often involve additional layers of complexity due to physiological and environmental variables such as growth stages, genotypic differences, environmental fluctuations (e.g., temperature, humidity, and photoperiod), seasonal variations, and soil composition, each potentially interacting uniquely with batch effects. For instance, dioecious plants (e.g., XX/XY system) may exhibit sex-specific responses to environmental batch conditions, complicating effective batch correction and emphasizing the need for biologically informed experimental designs (Dale et al. 2021, Upton et al. 2023). In recent years, DESeq2 (Love et al., 2014) has become one of the most widely used tools in RNA-seq analyses. It inherently accounts for batch effects directly within its modeling framework during differential expression analysis, making it unnecessary to correct count data before modeling. This approach has the advantage of preserving raw gene expression values, ensures the interpretability of biological signals and minimizes potential distortions associated with data transformation. However, in numerous analytical workflows — particularly those involving exploratory data visualization, clustering analyses, or dimensionality reduction techniques— batch-corrected expression values are desirable, as they significantly enhance interpretability and comparability among samples. Such downstream tasks often perform best with input data where batch-induced variations have already been mitigated.

To address this requirement, we present a novel batch correction method that leverages DESeq2’s internal model-based estimates to directly adjust raw count data, without relying on external correction tools or complex transformations. This approach preserves the integrity of log-fold changes between biological conditions and offers flexibility for handling complex experimental designs. Specifically, our approach effectively accommodates interaction effects, even in scenarios characterized by imbalanced or highly confounded settings. As a result, this facilitates improved data analysis and visualization, while maintaining compatibility with standard DESeq2 workflows.

### Implementation

The simulated dataset used in this study consists of 1,000 genes across 48 samples. The dataset is designed to be computationally efficient, with confirmed scalability to larger real-life datasets. The experimental design involves two batches, modeling gene expression in dioecious plants undergoing sex-specific treatments at two distinct time points: Day 0 (vegetative) and Day 1 (early flowering). Due to sex-specific treatments to alter flower sex, male and female plants were separated across batches: Batch 1 contained female controls (n=6) and treated males (n=6), while Batch 2 included male controls (n=6) and treated females (n=6). The treatments were applied following day 0 measurements but before day 1 measurements, allowing day 0 data to serve as a baseline for controlling batch effects. Samples (n=48) were collected across two genotypes, with three biological replicates per combination of sex, treatment, genotype and time point.

Two design models were implemented for batch correction analyses: a single-batch correction design (*~ sex + day + batch + sex:day + treatment*) and a double-batch correction design (*~ sex + day + batch + sex:day + sex:batch + treatment*). An additional, more complex model incorporating interacting factors (*~ sex + day + batch + genotype + sex:day + sex:batch + treatment + treatment:genotype*) was also designed.

Log_2_fold changes (LFC) calculated from DESeq2 were utilized to perform single and double-batch corrections, adjusting the expression levels to balance batch effects without biasing towards any specific batch.

For single-batch correction, original counts for gene *i*, with batches *A* and *B* and days *d* ∈ {0, 1}, are defined as 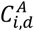 and 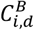. The DESeq2-estimated LFC between batches at day 0 is represented as *LFG*_*o*_. Adjusted counts for each time point, mitigating batch effects estimated at day 0, are calculated as:

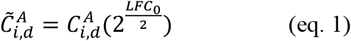

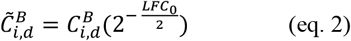

For double-batch correction, for any gene *i*, in male (*m*) or female (*f*) samples, batches labelled *A* and *B* and the days indexed as *d* ∈ {0, 1}, original (unadjusted) counts are defined as follows: 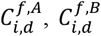 for female samples, and 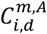 and 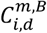 for male samples. DESeq2-estimated LFC between batches at day 0 are represented as 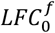 for females and 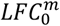 for males. To independently adjust counts for sexes at each time point, mitigating batch effects estimated at day 0, adjusted counts are calculated as:

Female samples:

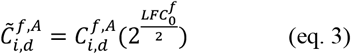

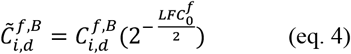

Male samples:

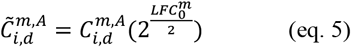

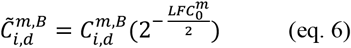

The GitHub repository (https://github.com/julienroyulaval/DESeq2-MultiBatch) provides detailed recommendations and practical strategies for performing DESeq2 analyses with complex experimental designs. It includes specific guidance on constructing design matrices, addressing imbalanced datasets, defining appropriate contrasts, and conducting multi-factorial analyses with illustrative examples.

Performance of batch correction method was benchmarked against limma’s removeBatchEffects (v. 3.58.1) (Ritchie et al. 2015) and ComBat-Seq (v. 3.50.0) (Zhang, Parmigiani and Johnson 2020) through principal components analysis (PCA) plots (ggplot2 version 3.5.1) (Wickham 2016) and heatmaps (pheatmap version 1.0.12) (Raivo Kolde 2010), in double-batch correction designs (Figure 1, Supplementary Figures 1). All analyses were performed in RStudio (v2024.9.0.375) running R (v4.3.1).

**Fig. 1:**
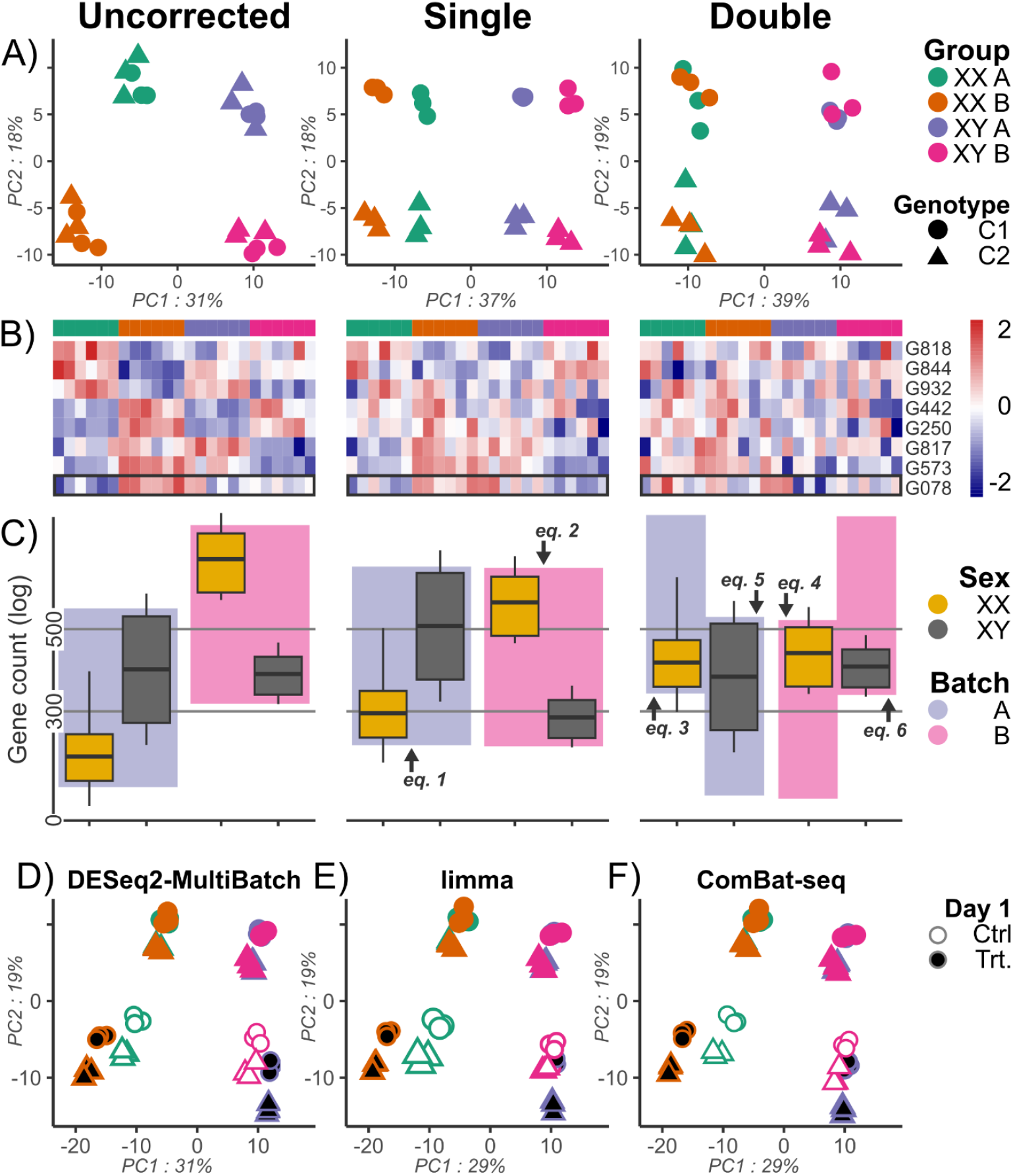
(A) PCA plots at day 0 of uncorrected, single batch corrected, and double batch corrected data using DESeq2-MultiBatch. (B) Heatmaps of nine highly batch-affected genes in uncorrected, single batch corrected, and double batch corrected data using DESeq2-MultiBatch, with Gene 078 framed. (C) Gene expression counts for G078 in uncorrected, single batch corrected, and double batch corrected data using DESeq2-MultiBatch, (eq.1-6) shown as arrows. (D-F) PCA plots comparing batch correction (day 0 and day 1) using DESeq2-MultiBatch, limma’s removeBatchEffects, and ComBat-seq.

## Results

Analyses of the simulated dataset from dioecious plants showed substantial batch in uncorrected data at day 0 (Figure 1A, Supplementary Figure 2 for all uncorrected data), primarily resulting from the experimental design. Application of single batch correction (eq. 1 and eq. 2) significantly reduced batch-induced variation; however, residual differences among biologically identical groups remained evident, as demonstrated by distinct clusters, highlighting persistent *batch* and *sex* interactions (Figure 1A). In contrast, the sex-dependent batch-correction method, explicitly accounting for these interactions, significantly improved the clustering of biologically identical groups, effectively removing batch-associated variation while preserving inherent biological variability within-groups (Figure 1A). Post-correction analysis confirmed a complete elimination of differentially expressed genes between batches at day 0, separately for both males and females.

Heatmap analyses of eight highly batch-affected genes further quantified correction effectiveness. Prior to any correction, clear batch-related expression patterns were observed (Figure 1B). Single batch correction partially reduced this variation but did not eliminate discrepancies among same-sex groups, for example in G078, which is highly affected by *sex* and *batch* interactions. Conversely, sex-dependent correction successfully normalized expression levels across batches for all selected genes, substantially reducing batch-induced variations.

Gene expression counts of G078 (Figure 1C) show a notable discrepancy in female samples between batches A and B prior to correction, which is not seen in male samples. Correcting for a homogeneous batch effect skews the values for male samples. The sex-dependent correction method independently adjusts male and female expression levels, leading to a more precise alignment of the gene expression data between batches for each sex separately. This approach proved especially advantageous for genes whose expression levels were influenced by interaction between sex and environmental factors across batch.

Benchmarking our method against established batch correction approaches shows comparable or superior performance in reducing experimental batch effects. PCA plots encompassing both time points (day 0 and day 1) showed that sex-dependent batch correction provided tighter clustering and clearer separation of biological groups (Figures 1D), closely aligning with results obtained from standard methods such as limma’s removeBatchEffects (Figures 1E, Supplementary Figure 1 for heatmap) and ComBat-seq (Figures 1F, Supplementary Figure 1 for heatmap).

RNA-seq has become a fundamental technique in biological research across diverse species and experimental systems; nonetheless, analyses frequently encounter challenges due to batch effects, which obscure genuine biological signals. Although most literature on batch-effect correction algorithms primarily addresses technical batch effects originating from sample preparation, sequencing, or processing steps, complex and multifactorial studies require effective strategies to correct experimental batch effects that may involve interaction factors.

Our proposed DESeq2-MultiBatch method effectively addresses this issue by explicitly modeling interaction effects within the DESeq2 analytical framework, significantly enhancing sample clustering and alignment while preserving genuine biological variability. Compared to alternative methods such as limma’s removeBatchEffect and ComBat-seq, DESeq2-MultiBatch provides a more intuitive workflow by directly modeling interaction effects without requiring complex relabeling of metadata. Additionally, unlike limma’s removeBatchEffect, which operates on variance-stabilized data, DESeq2-MultiBatch retains the ability to produce batch-corrected expression counts directly from raw data, enhancing interpretability and downstream utility.

Despite its strengths, the efficacy of DESeq2-MultiBatch depends critically on accurate initial modeling of experimental conditions. Misspecification or inadequate representation of interaction factors may limit correction effectiveness and risk attenuating true biological variation. Looking forward, integrating DESeq2-MultiBatch into widely used RNA-seq analytical pipelines presents a promising avenue for enhancing the robustness and accuracy of multi-factorial RNA-seq experiments. Overall, DESeq2-MultiBatch represents a practical, accessible, and robust solution to the pervasive challenge of batch effect correction in RNA-seq analyses.

## Supporting Information

