## Supplementary figures for "DESeq2-MultiBatch: Batch Correction for Multi-Factorial RNA-seq Experiments"

­**Supplementary material**

**Figures**

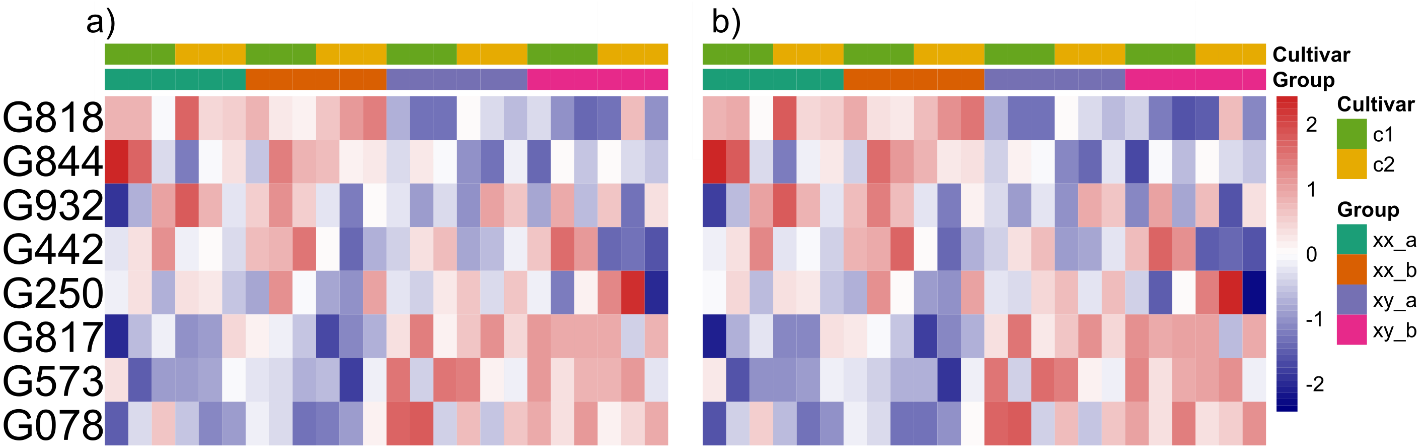

Supplementary Figure 1: a) Heatmap of eight highly batch-affected genes after limma’s removeBatchEffects. b) Heatmap of eight highly batch-affected genes after ComBat_seq.

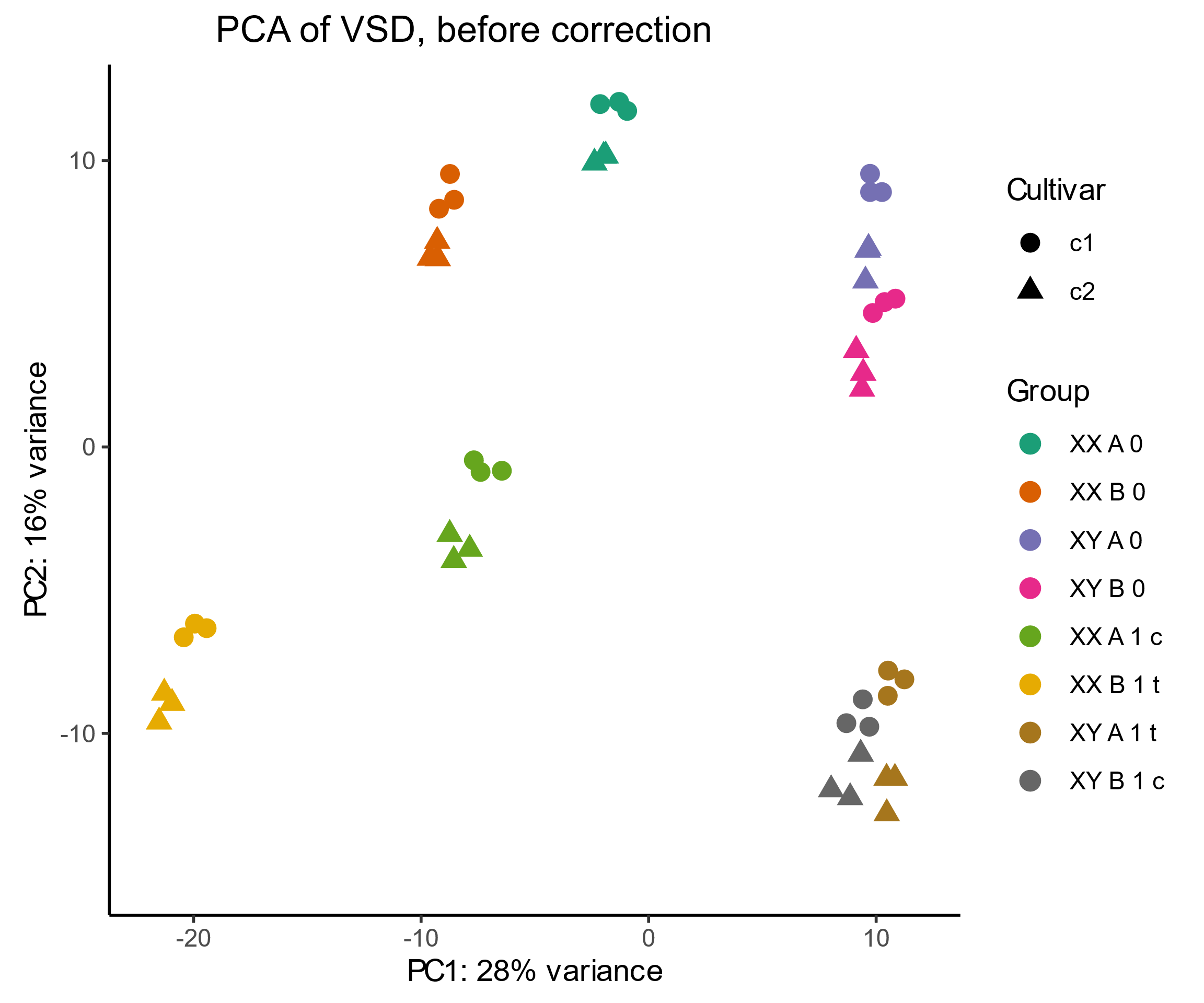

Supplementary Figure 2: PCA plot of uncorrected data for all samples.
